# Accurate prediction of residue-residue contacts across homo-oligomeric protein interfaces through deep leaning

**DOI:** 10.1101/2020.09.13.295196

**Authors:** Yumeng Yan, Sheng-You Huang

## Abstract

Protein-protein interactions play a fundamental role in all cellular processes. Therefore, determining the structure of protein-protein complexes is crucial to understand their molecular mechanisms and develop drugs targeting the protein-protein interactions. Recently, deep learning has led to a breakthrough in intraprotein contact prediction, achieving an unusual high accuracy in recent CASP structure prediction challenges. However, due to the limited number of known homologous protein-protein interactions and the challenge to generate joint multiple sequence alignments (MSA) of two interacting proteins, the advances in inter-protein contact prediction remain limited. Here, we have proposed a deep learning model to predict inter-protein residue-residue contacts across homo-oligomeric protein interfaces, named as DeepHomo, by integrating evolutionary coupling, sequence conservation, distance map, docking pattern, and physic-chemical information of monomers. DeepHomo was extensively tested on both experimentally determined structures and realistic CASP-CAPRI targets. It was shown that DeepHomo achieved a high accuracy of >60% for the top predicted contact and outperformed state-of-the-art direct-coupling analysis (DCA) and machine learning (ML)-based approaches. Integrating predicted contacts into protein docking with blindly predicted monomer structures also significantly improved the docking accuracy. The present study demonstrated the success of DeepHomo in inter-protein contact prediction. It is anticipated that DeepHomo will have a far-reaching implication in the inter-protein contact and structure prediction for protein-protein interactions.

## 1 INTRODUCTION

As one of the most important biological macromolecules in living organisms, proteins have evolved to conduct diverse cellular functions, from serving as reaction catalysts to regulating DNA translation and transcription. Proteins rarely work in monomeric form but are rather assembled in homooligomers or hetero-oligomers to perform their biological functions [1, 2]. The structural characterization of protein-protein interactions and high-order assemblies is therefore crucial to elucidate the molecular mechanisms behind them and understand the related biological processes [3, 4]. Despite the great progress in experimental determination of macromolecular structures [5–7], the number of protein structures that have been experimentally determined and deposited in the Protein Data Bank (PDB) [8] is still limited. Compared with the limited structures in the PDB, the genetic sequential information has increased dramatically as the result of high-throughput sequencing technologies and large-scale genome projects [9, 10]. Taking advantage of the huge sequence information through sophisticated statistical models provides a valuable alternative to determine the structures. Recently, direct coupling analysis (DCA) [11–15] and similar tools [16–19], which try to distinguish direct from indirect correlation effects, have been developed to improve the performance of residue-residue contact prediction from multiple sequence alignments (MSAs). Such computational methods have proven to be very useful in monomer protein structure prediction in the Critical Assessment of protein Structure Prediction (CASP) experiments [20–23].

Subsequently, such DCA methods have been extended from the contact prediction in monomer protein structure to residue-residue prediction in hetero protein-protein interface [24, 25]. Despite some successes, there is still a challenge for such coevolution-based contact prediction in proteinprotein interactions. That is, how to create a joint MSA of high quality, in which each line contains the sequences of a pair of interacting proteins [26]. Constructing an MSA for individual proteins is relatively easy using protein sequence searching software like HHblits [27] or PSI-blast [28]. However, generation of the joint MSA from the two MSAs of interacting proteins is difficult due to the existence of paralogs [26, 29–31]. In bacteria, the two individual MSAs can be concatenated according to their genomic distance because the genes coding for interacting proteins are often colocalized in the same operon [24, 25]. For eukaryotes, phylogeny tree may be used to concatenate MSAs [32].

Recently, two iterative methods have been presented to identify specific interacting paralogs by maximizing the DCA signals [29, 30]. However, all above methods have limitations because the produced MSA does not have sufficiently divergent sequences or contains many incorrect paired sequences. Besides the DCA models, some machine/deep learning methods have been developed to predict the reside-residue contacts between protein-protein interfaces with [32] or without producing the joint MSA for the complex [33–36]. However, the accuracy of these methods is still relatively low because of the impact of joint MSAs.

To avoid the joint MSA challenge, Uguzzoni *et al.* have focused on the interactions between homo-oligomers rather than hetero-oligomers, for which the produced individual MSA can be directly used to perform the statistical analysis on the basis of the assumption that the sequences in each line of the MSA also form homo-oligomer interfaces like the queried proteins [37]. This avoids the challenge of creating of the joint MSA for residue-residue contact prediction between hetero-oligomeric protein interfaces. However, another problem presents, that is how to distinguish the intraprotein and interprotein contacts in a homo-oligomer. Uguzzoni *et al.* assumed that the predicted residue pair contacts with long intramonomer distances are inter-protein contacts. However, using the monomer structure to filter all the intra-protein contacts may neglect the residue-residue contacts that are both intra-protein and inter-protein.

Recently, deep learning has achieved great successes in intraprotein residue-residue contact prediction and demonstrated high accuracy in the 12th and 13th Critical Assessment of protein Structure Prediction (CASP12 and CASP13) structure prediction challenges [38–45]. However, there are significant differences between monomer proteins and homo-oligomer complexes in terms of contact prediction. First, for homo-oligomer interface contact prediction, it is difficult to remove the potential intra-protein contact noises which exist in the MSA. Second, the number of inter-protein contacts is far smaller than that of intra-protein contacts for homo-oligomers. This means that the positive and negative labels are much more unbalanced in the inter-protein contact prediction for homo-oligomers than the intra-protein contact prediction for monomer proteins. To overcome these challenges, we have here presented a deep learning model to predict residue-residue contacts across homo-oligomer interfaces, named as DeepHomo, by integrating evolutionary coupling, sequence conservation, structural features, and physic-chemical information of monomers. We have used Focal Loss function [46] to focus learning on the less hard examples to deal with the unbalance between the numbers of contacted and non-contacted residue pairs. DeepHomo was extensively tested on 300 diverse homo-oligomer proteins from the PDB and 28 realistic targets from recent Critical Assessment of protein Structure Prediction-Critical Assessment of Predicted Interactions (CASP-CAPRI) challenges [47–49]. It was shown that DeepHomo outperformed stat-of-the-art DCA and machine learning-based approaches and led to a significant improvement in the docking accuracy for structure prediction of homo-oligomers.

## 2 RESULTS

### 2.1 Workflow of DeepHomo

Figure 1 shows the workflow of our deep learning-based inter-protein contact prediction for homo-oligomer complexes (DeepHomo). DeepHomo starts with a monomer structure that may be taken from an experimental structure or predicted by a third-party protein structure prediction method like I-TASSER [50, 51]. DeepHomo is designed to take full advantage of both the structure and sequence information of monomers, which can also be grouped into 1D sequential and 2D pairwise features. One one hand, the 1D sequential features, including the secondary structure (SS), hydrophobicity, and position-specific scoring matrix (PSSM) information, are first extracted from the monomer structure/sequence. Next, the 1D ResNet CNN is used to learn the high-dimensional features from the 1D sequential features. Then, a 2D pairwise matrix is constructed by outer concatenation from the high-dimensional sequential features. On the other hand, the 2D features, including the intra-residue distance map, docking map, and direct coupling analysis (DCA) scores are obtained from the structure and MSA of the monomer. Then, these 2D matrices plus the previously converted 2D map from sequential features are fed into a 2D ResNet CNN for training, resulting in the final matrix of predicted contacts (Figure 1).

**Figure 1:**
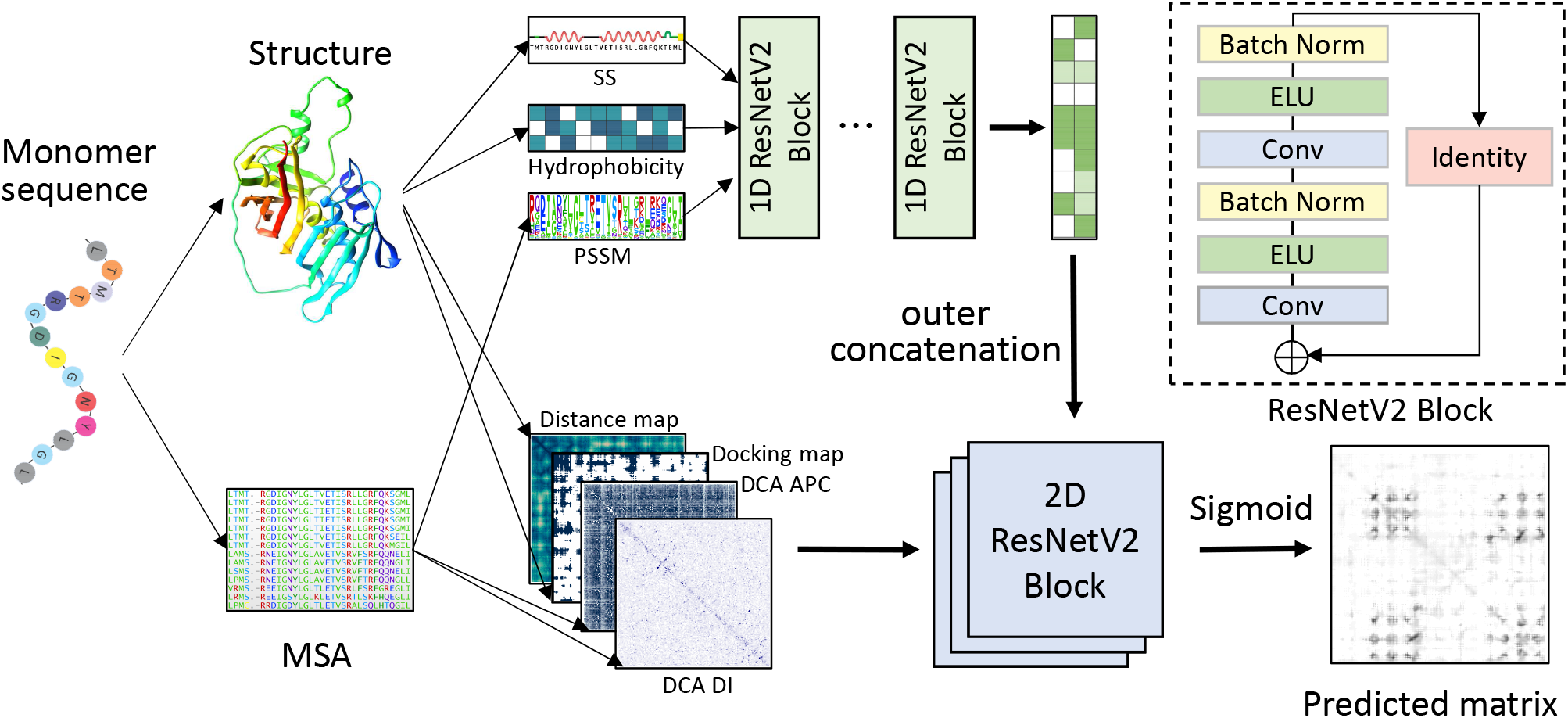
The framework of the DeepHomo model. The model consists of two components: a 1D convolutional neural network with 1D features as input which contain secondary structure (SS), hydrophobicity features, and PSSM matrix (top) and a 2D convolutional neural network (bottom) with 2D features as input which include distance map, docking map and DCA scores.

### 2.2 Evaluation on homo-dimeric structures from the PDB

#### 2.2.1 Performance of DeepHomo

We first evaluated the performance of our DeepHomo model in inter-protein residue-residue contact prediction on a large test set of 300 experimental homo-dimer complexes from the PDB. Figure 2 shows the accuracy and success rate of DeepHomo in residue contact prediction as a function of the number of predicted contacts on the 300 homo-dimers. For comparison, the figure also shows the corresponding results of direct coupling analysis (DCA)-based methods on this test set. The corresponding results for top 1, 10 and 100 predicted contacts are listed in Table 1. The DCA-based methods used as baselines in this study is similar to the Uguzzoni *et al*’s method [37]. First, based on the produced MSA, the direct coupling scores containing the raw direct information (DI) score and APC-corrected score were calculated using CCMpred. Then, according to the input monomer structure, the residue pairs with intra-monomer distances of more than 12 Å were regarded as the candidates of inter-protein contacts for the homo-oligomeric interface. At last, the candidate contacts were sorted by their co-evolutional scores, DI scores or APC scores, which represent DCA_DI and DCA_APC contact prediction approaches. It can be seen from Table 1 and Figure 2 that our DeepHomo model achieved a much better performance than DCA-based methods. For the top 1, 10 and 100 predicted contacts, DeepHomo obtained the accuracies of 62.3%, 52.6% and 37.6%, respectively, compared with 27.0%, 15.6% and 5.2% for DCA_DI and 33.0%, 22.2% and 8.1% for DCA_APC. Similar advantages of DeepHomo over DCA-based approaches can also be observed in the success rate of contact prediction. For example, DeepHomo gave a high success rate of 94% for top 100 predicted contacts, which is significantly higher than 75.0% for DCA_APC and 65.7% for DCA_DI (Table 1 and Figure 2b). Although our deep learning model used the same MSA and monomer structure with DCA-based methods, the accuracy of DeepHomo almost doubled the accuracy of DCA_APC when the top predicted contact was considered. For top 100 predicted contacts, the improvement is much more significant and the accuracy has been improved almost over four times, compared with DCA_APC method. For DCA-based methods, DCA_APC method performed better than DCA_DI method for all the top 100 predicted contacts. This is consistent with the previous study in monomer protein contact prediction that average product correction helps improve the prediction quality [15, 52].

**Table 1:**
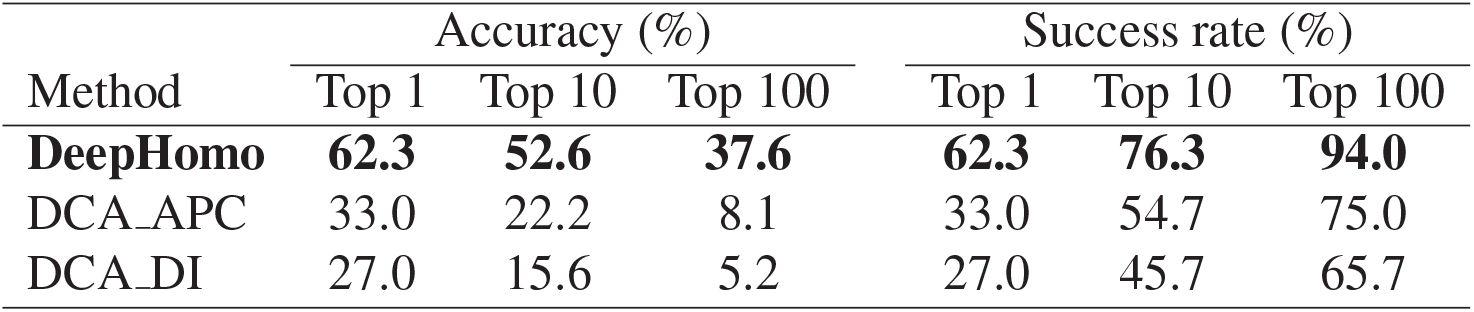
Comparison of the results by DeepHomo and two DCA-based methods on the PDB test set of 300 experimental homo-dimer complexes when the top 1, 10 and 100 predicted contacts are considered.

| Method | Accuracy (%) |  |  | Success rate (%) |  |  |
| --- | --- | --- | --- | --- | --- | --- |
|  | Top 1 | Top 10 | Top 100 | Top 1 | Top 10 | Top 100 |
| **DeepHomo** | **62.3** | **52.6** | **37.6** | **62.3** | **76.3** | **94.0** |
| DCA_APC | 33.0 | 22.2 | 8.1 | 33.0 | 54.7 | 75.0 |
| DCA_DI | 27.0 | 15.6 | 5.2 | 27.0 | 45.7 | 65.7 |

**Figure 2:**
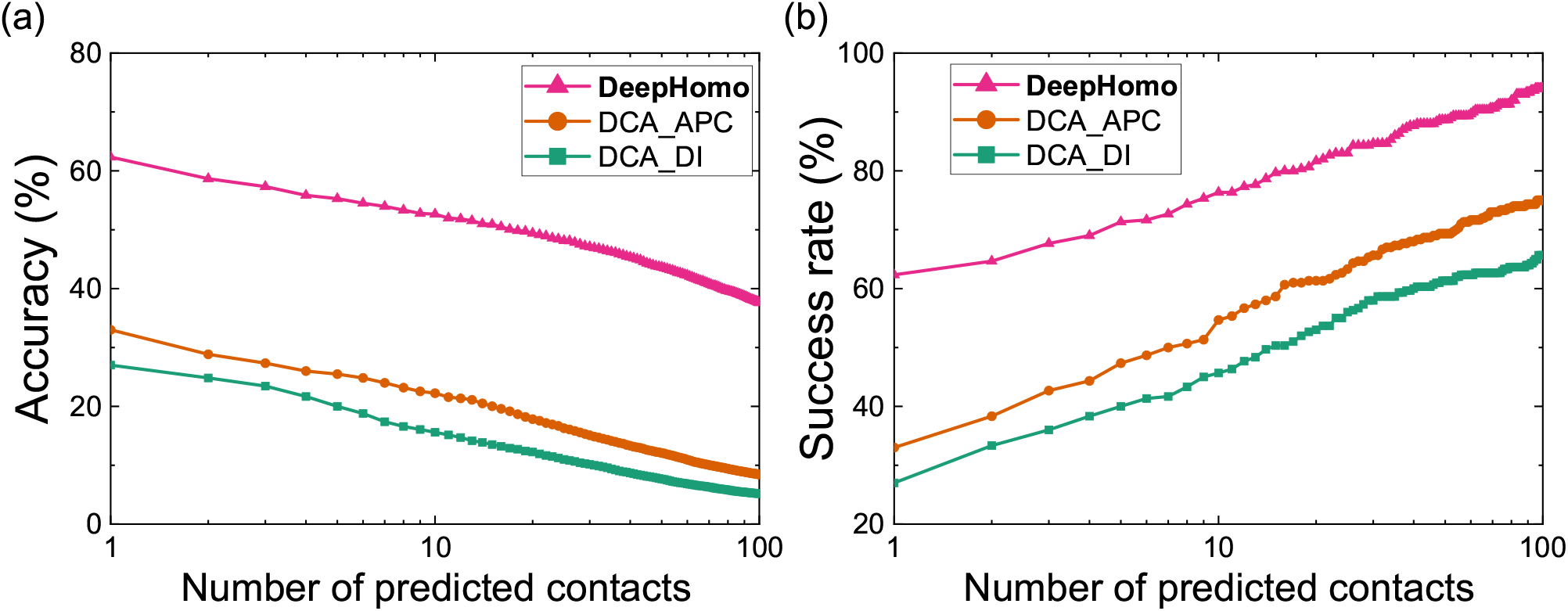
The performance of DeepHomo and two DCA-based approaches (DCA_DI and DCA_APC) on the PDB test set of 300 experimental homo-dimeric structures. (a) The accuracy, number of TP predictions divided by number of predictions, as a function of the number of predicted contacts. (b) The success rate, number of the targets with at least one successfully predicted contact divided by the total number of targets in a test set, as a function of the number of predicted contacts.

#### 2.2.2 Impact of different features

To investigate the effect of different features on the performance of DeepHomo, we have conducted the ablation studies on the PDB test set of 300 homo-dimers. In the ablations studies, all the hyper-parameters were fixed, only the input features were changed during each experiment. Our DeepHomo model has six types of features, including distance map of the monomer structure, DCA scores, docking map, secondary structure, PSSM matrix and hydrophobicity features. We first investigated the influence of each of the six features on the performance of our model by excluding that feature from the input. Figure 3a shows the accuracies of our model for top 1 and 10 predicted contacts after excluding different features from the input. It can be seen from the figure that all six features have impacts on the performance of DeepHomo as removing any feature will result in some degree of decrease in the accuracy. Among the six features, the DCA-based feature is the most important feature, because without the feature, the accuracy shows the largest decrease from 62.3 and 52.6% to 49.3 and 42.5% for top 1 and 10 predicted contacts (Figure 3a). This can be understood because the DCA information has also been found to be the important feature in intra-protein contact prediction [41]. Removing the hydrophobicity features and PSSM matrix from input has little effect on the top 10 accuracy of our model. It can be explained that these two features have overlap with other features to some extent, so the information encoded in these two features can also be extracted from the other features by the convolutional neural network. Similarly, the secondary structure feature may have the similar information with the hydrophobicity features. The DCA scores may encode the similar evolutionary information with the PSSM matrix as they are both derived from the same MSA.

**Figure 3:**
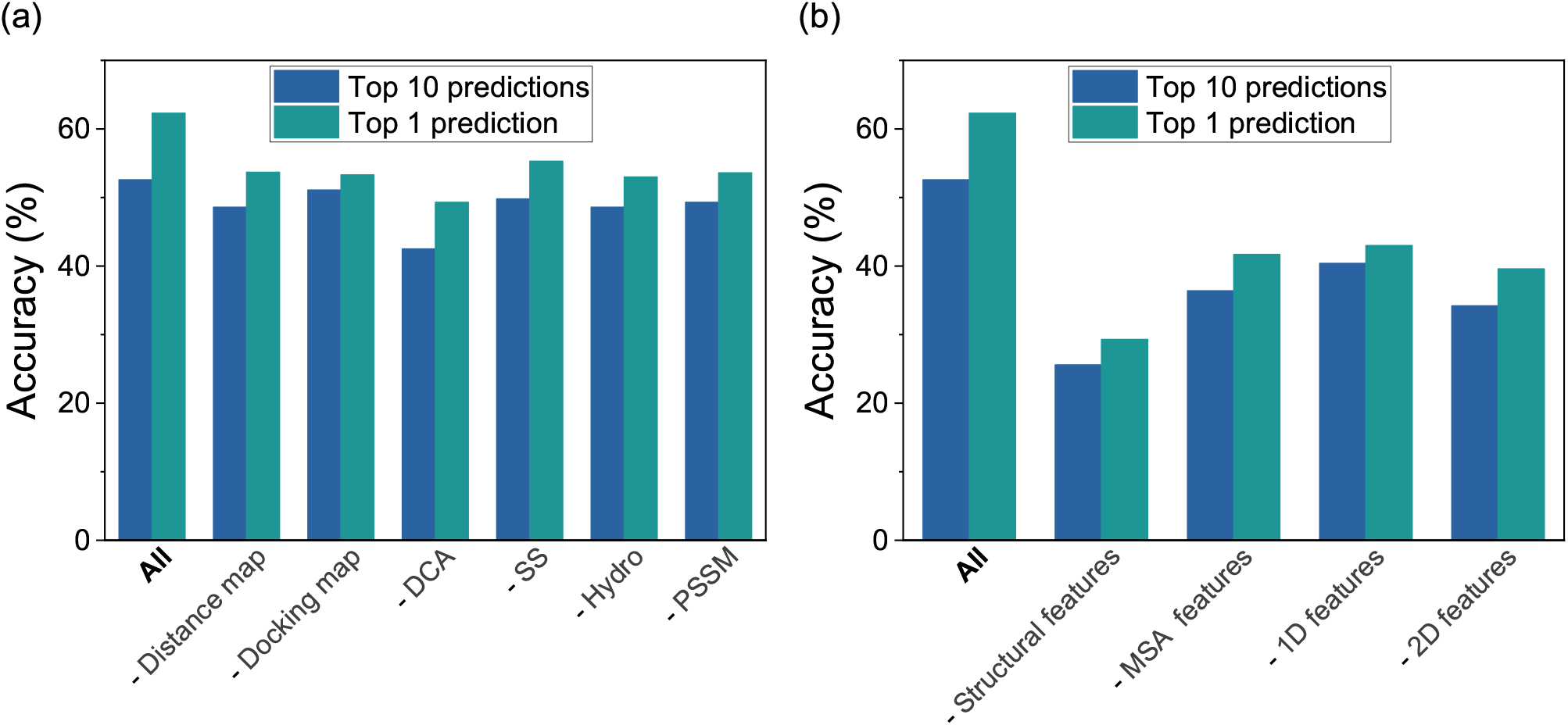
Ablation experiments of DeepHomo on the PDB test set. ‘All’ stands for the original DeepHomo model using all the features. Each label starting with “-” represents the experiment after excluding the corresponding feature(s). (a) Ablation results of six individual features. (b) Ablation results of four types of features.

According to the dimension of input features, these features can also be divided into 1D features which include secondary structures, PSSM matrix, hydrophobicity features, and 2D features which including distance map, DCA scores and docking map. From the results of ablation experiments shown in Figure 3b, we can see that both 1D and 2D features are important because eliminating any of these features resulted in a significant decrease in the accuracy. To investigate the influence of the monomer structure and the MSA, we also did the ablation study for structural features which include distance map, docking map, secondary structure and two hydrophobic features, and MSA features which include DCA scores and PSSM matrix. As shown in Figure 3b, the structural features and MSA features are both very important to our model. It is obviously that the MSA features is critical to our deep learning model where the information of spatial adjacencies between residues may be hidden in the evolutionary information. Compared with the MSA features, the structural features have a higher impact on the accuracy of DeepHomo (Figure3b). This is interesting because the present finding means that the complex structure between two proteins depends on the structure more than the sequence of monomers. In other words, those monomers with structural analogy tend to form similar complex structures than those with sequence homology.

#### 2.2.3 Impact of the MSA depth

As the MSA is crucial to our DeepHomo model and other DCA-based approaches, we have examined the effect of the MSA depth on the performance of our method. Here, the effective number of sequence homologs in an MSA, *M*_eff_, was used to measure the MSA depth. *M*_eff_ can be regarded as the number of non-redundant sequence homologs in an MSA under a sequence identity cutoff. The sequence identity between two sequences in an aligned MSA is defined as the percentage of the positions with corresponding residues. In this study, a sequence identity cutoff of 70% was used to remove redundancy. The *M*_eff_ is calculated as follows:

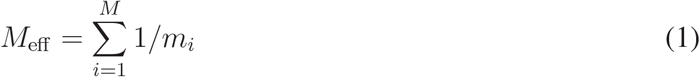

where *M* is the number of sequences in an MSA and *m*_*i*_ is the number of sequences that have sequence identity >70% with the *i*-th sequence in the MSA.

Figure 4 shows that the accuracies of top 1 and top 10 predicted contacts with respect to ln(*M*_eff_) on the test set. The first 3 intervals were merged together because of the small number of proteins in these intervals. It can be seen from the figure that our DeepHomo method outperformed the two DCA-based approaches regardless of the number of *M*_eff_. As the evolutionary information is included in the MSA, all methods obtained a better performance with increased *M*_eff_ for both top 1 and top 10 predicted contacts. A higher *M*_eff_ generally leads to better coevolutionary scores for the DCA-based methods and evolutionary input features for our deep learning model, which resulted in a better performance of all tested methods. In addition, despite the high accuracy for the cases of ln(*M*_eff_) > 6 (approximately *M*_eff_ > 400), our DeepHomo model also obtained good accuracies in the 4-5 and 5-6 ranges of ln(*M*_eff_), and gave the accuracies of 58.3% and 58.3% for top 1 predicted contact and 43.3% and 40.8% for top 10 predicted contacts, respectively. This suggests that DeepHomo is able to learn correct contact information from not very deep MSAs with the help of other information such as structural features.

**Figure 4:**
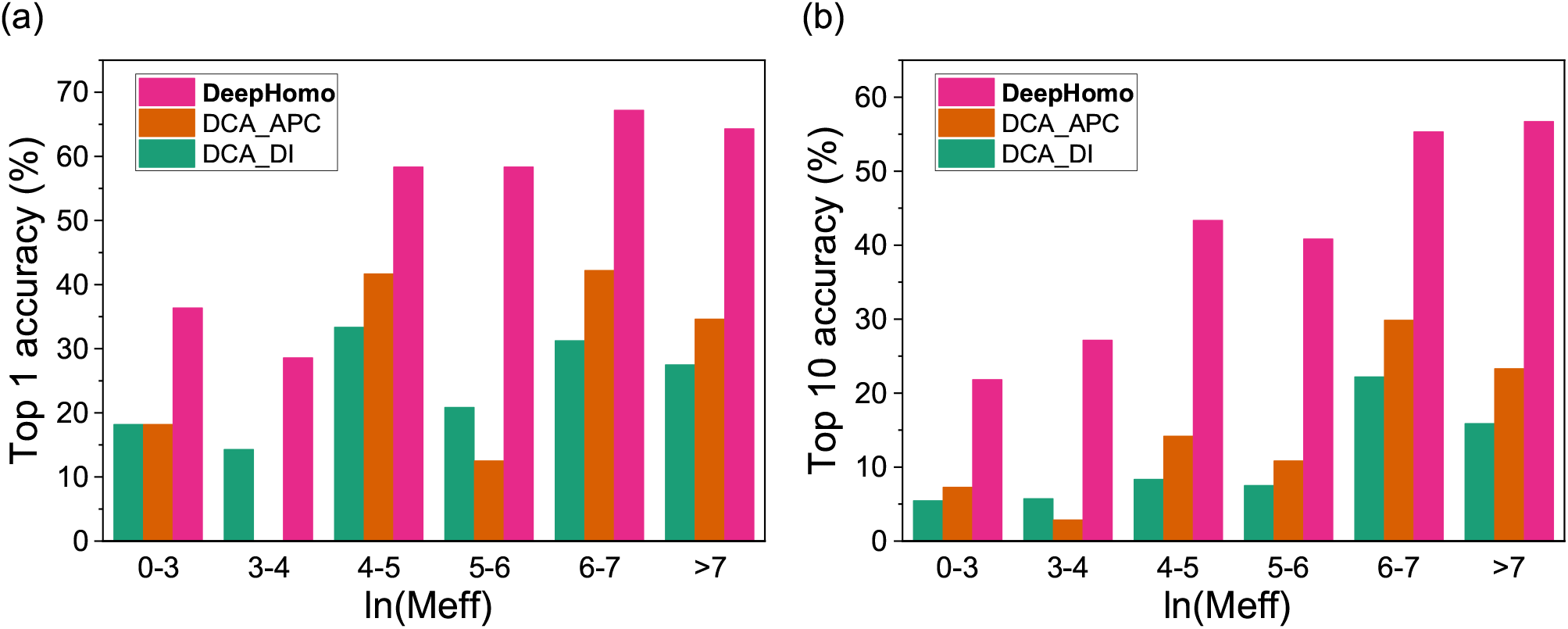
The accuracies of DeepHomo, DCA_DI and DCA_APC with respect to the MSA depth measured by ln(*M*_eff_) on the PDB test set of 300 homo-dimeric structures. (a) The results for the top predicted contact. (b) The results for the top 10 predicted contacts.

### 2.3 Application to realistic CASP-CAPRI targets

#### 2.3.1 Performance of DeepHomo

As the monomer structure for a sequence is normally unknown in real applications, we have further tested our DeepHomo model on the CASP-CAPRI set of 28 realistic homo-oligomer targets. To be consistent with the blind prediction nature in CASP, we have used the top Zhang-Server model, which has been blindly predicted by the I-TASSRER server from the Zhang group [50, 51], as the monomer structure for the input of DeepHomo during the evaluations.

Figure 5 shows the accuracy and success rate of DeepHomo in residue contact prediction as a function of the number of predicted contacts on the CASP-CAPRI test set. For comparison, the figure also gives the corresponding results of other DCA-based methods and machine learning-based BIPSPI approaches. The accuracies and success rates for top 1, 10 and 100 contacts are listed in Table 2. BIPSPI is a machine-learning method for the prediction of residue-residue contacts in the hetero-dimer interfaces which employs Extreme Gradient Boosting (XGBoost) models as a classifier [36]. It was trained on the Protein-Protein Docking Benchmark version 5.0 [53] and has achieved a good performance in the test. BIPSPI has two versions that accepts the sequence as input (BIPSPI_seq) or the structure as input (BIPSPI_struc), respectively. We have tested both the sequential and structural versions by submitting jobs to the BIPSPI web server. The monomer structure uploaded to the web server is the same as that used by our DeepHomo model. Once the jobs done, the predicted results were downloaded from the BIPSPI web server and analyzed.

**Table 2:** Comparison the results by DeepHomo and other four approaches on the CASP-CAPRI set of 28 realistic targets when the top 1, 10 and 100 predicted contacts are considered.

| Method | Accuracy (%) |  |  | Success rate (%) |  |  |
| --- | --- | --- | --- | --- | --- | --- |
|  | Top 1 | Top 10 | Top 100 | Top 1 | Top 10 | Top 100 |
| **DeepHomo** | **60.7** | **52.5** | **34.3** | **60.7** | **67.9** | **85.7** |
| DCA_APC | 14.3 | 17.5 | 7.3 | 14.3 | 42.9 | 75.0 |
| DCA_DI | 25.0 | 12.1 | 3.9 | 25.0 | 32.1 | 53.6 |
| BIPSPI_seq | 7.1 | 5.4 | 5.1 | 7.1 | 25.0 | 75.0 |
| BIPSPI_struc | 25.0 | 14.6 | 14.5 | 25.0 | 57.1 | 85.7 |

**Figure 5:**
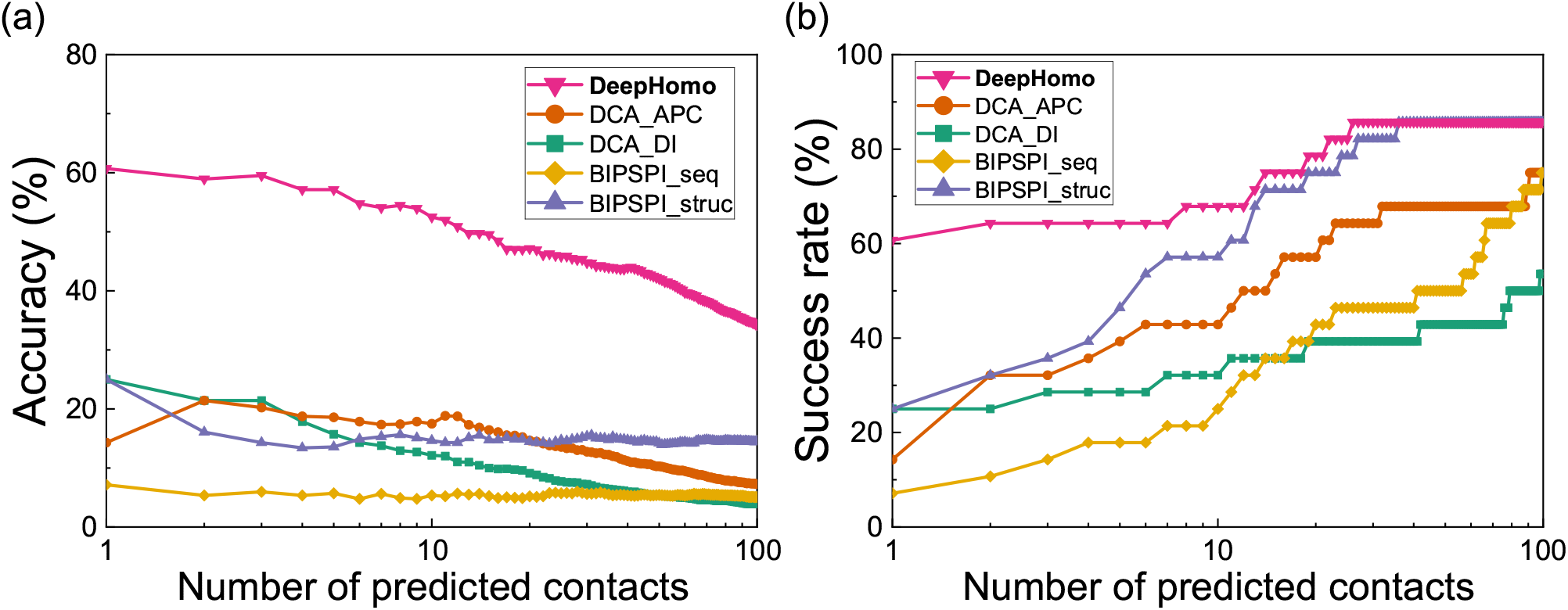
The performance of DeepHomo, two DCA-based approaches (DCA_DI and DCA_APC) and two machine learning-based methods (BIPSPI_seq and BIPSPI_struc) on the CASP-CAPRI of 28 realistic homo-dimeric targets. (a) The accuracy as a function of the number of predicted contacts. (b) The success rate as a function of the number of predicted contacts.

It can be seen from Figure 5 that DeepHomo achieved the best performance among the five methods and gave a much higher accuracy and success rate than the other four methods. Specifically, DeepHomo obtained the accuracies of 60.7%, 52.5% and 34.3% for top 1, 10 and 100 predicted contacts, while the second-best method BIPSPI_struc gave much lower accuracies of 25.0%, 14.6% and 14.5%, respectively (Table 2). Similar trends can also be observed in the success rates of different approaches, showing that DeepHomo yielded an overall much better performance than the other four approaches. Another notable feature in Figure 5b is that among the four approaches except Deep-Homo, BIPSPI_struc obtained an overall better performance than the rest three approaches. Given the importance of structural features in contact prediction, this finding can be understood because BIPSPI_struc used the monomer structure as input in its contact prediction, while the other three approaches only took the sequence as input. Comparing Figures 2a and 5a, one can also find that DeepHomo maintained a comparably high accuracy of >60% on both the PDB test set of 300 homo-dimers and the CASP-CAPRI test set of 28 realistic targets. Given that the PDB test set consists of experimental monomer structures while the CASP-CAPRI test set is formed by blindly predicted models, the similar accuracy of DeepHomo on the two test sets suggested that DeepHomo is robust to the quality of monomer structures. This is especially valuable because the monomer structure is often unknown and needs to be predicted by a structure prediction method in realistic applications.

#### 2.3.2 Impact of monomer structural quality

To investigate the impact of the quality of monomer structures, we have examined the performance of our DeepHomo model on the monomer structures with different accuracies in the CASP-CAPRI test set. Here, TM-score [54] was used to measure the quality of a monomer structure. For each target in the CASP-CAPRI test set, the TM-score between the predicted monomer structure and the native structure was calculated using TM-align [55]. Figure 6 gives the average accuracies of DeepHomo for top 1 and top 10 predicted contacts in the different bins of TM-score. For comparison, the figure also shows the corresponding results of other three approaches, DCA_DI, DCA_APC, and BIPSPI_struc. Here, BIPSPI_seq was excluded because it does not need monomer structures in the contact prediction. In addition, as no TM-score is in the range 0.5-0.6, no result is shown for this interval in the figure. All targets with TM-score<0.5 were merged in one interval [56].

**Figure 6:**
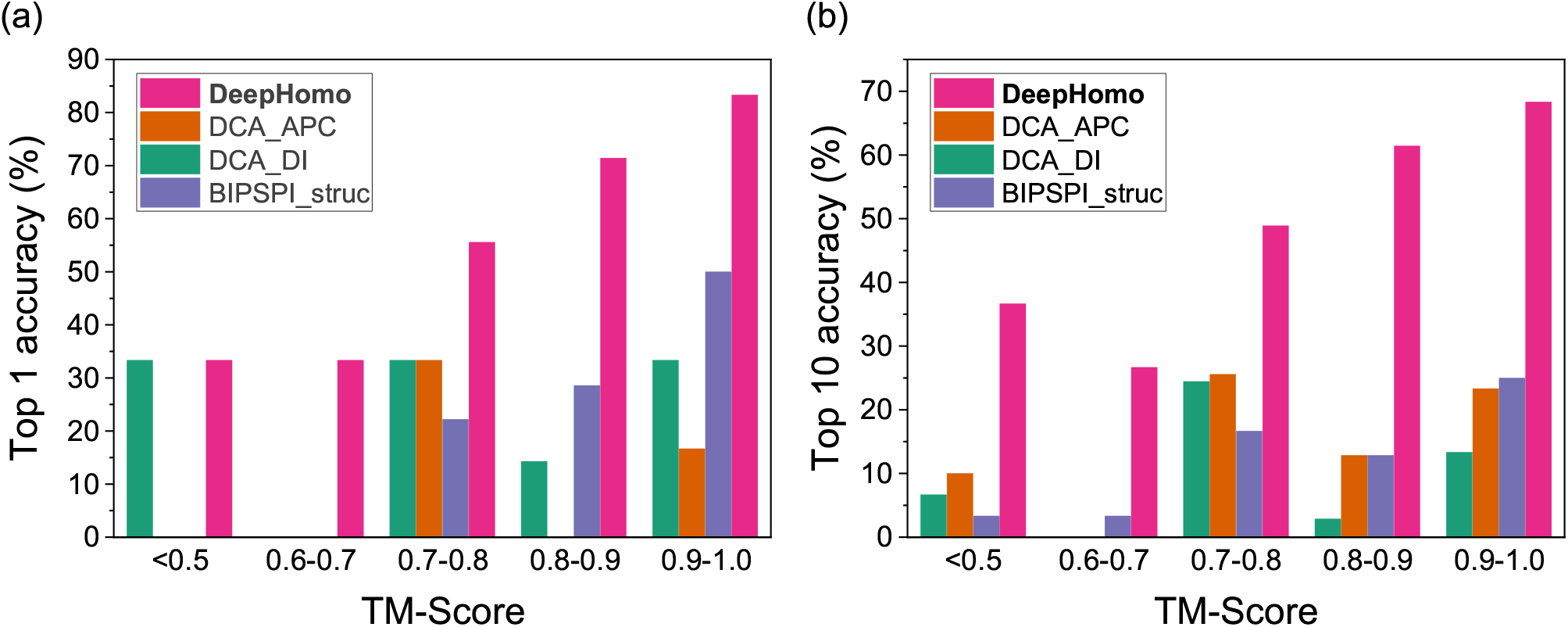
The accuracies of the top 1 (a) and top 10 (b) predicted contacts by DeepHomo and other four methods with respect to the quality of monomer structures measured by TM-score on the CASP-CAPRI test set of 28 realistic targets.

From Figure 6, one can see a general trend that DeepHomo and BIPSPI_struc show better accuracies for better-quality monomer structures with higher TM-scores, while the performance trend is not very clear for the DCA-based methods. It can also be seen from Figure 6 that the TM-score of 0.7 seems to be a threshold for different approaches. On one hand, the accuracies of the DCA-based methods for TM-score>0.7 become much better than those for TM-score <0.7 when the top 10 predicted contacts were considered. This may be understood because the targets with TM-score <0.7 tend to be hard targets and do not have many homologous sequences. Therefore, the DCA-based approaches would not be able to extract significant coupling information due to the limited number of sequences in the MSA. On the other hand, DeepHomo performed better with the increased quality of monomer structures for TM-score>0.7, and reached a stable accuracy of about 32% for TM-score<0.7 when the top predicted contact was considered. From Figure 6, one can also see that overall DeepHomo and BIPSPI tend to depend on the TM-score of monomer structures more than the DCA-based approaches. This can be understood because both DeepHomo and BIPSPI take structural features extracted from the monomer structure as input. However, for the DCA-based methods, the monomer structure was only used as a filter in the post process, so these methods rely less on the quality of the monomer structures. Despite the impact of structural quality, our DeepHomo model can achieve a good accuracy of 55.5% for targets with TM-score in 0.7-0.8 for the top predicted contact. This means that our deep learning model is accurate enough to learn correct contact information with only moderate-quality monomer structures.

#### 2.3.3 Examples of contact prediction

Figure 7 shows two selected examples of the top 100 predicted contacts by our DeepHomo model and the DCA_APC method versus the native contact map for targets T85 and T93 in the CASP-CAPRI test set. It can be seen from the figure that the predicted contacts by our DeepHomo model are all distributed near the native ones and can grab the important interaction mode. However, the predicted contacts by the DCA_APC method are very dispersed across all the contact map, which results in many false positive predicted contacts. Specifically, DeepHomo achieved the high accuracies of 69% and 84% on the two targets of T85 and T93, respectively, while the accuracies of the DCA_APC method were only 25% and 31%. From Figure 7, one can also see that the predicted contacts by DeepHomo are mostly close to the native contacts on the contact map, even though they may not overlap. That means that such near-native contacts by DeepHomo are still roughly correct, even though they may be classified to be incorrect contacts according the cutoff of 8 Å. In contrast, for the DCA_APC method, many of its predicted contacts are far from the native ones and thus are truly wrong contacts (Figure 7). Figure 8 shows a comparison of the top 10 predicted contacts by DeepHomo and DCA_APC in the native homo-dimeric structures of T85 and T93. It can be seen from the figure that our DeepHomo model successfully predicted all the contacts and achieved an accuracy of 100% on Targets T85 and T93 for top 10 predicted contacts, while the DCA_APC method only gave correct predictions for 20% and 80% of the native contacts on these two targets (Figure 8). These results again suggest the accuracy and robustness of our DeepHomo approach.

**Figure 7:**
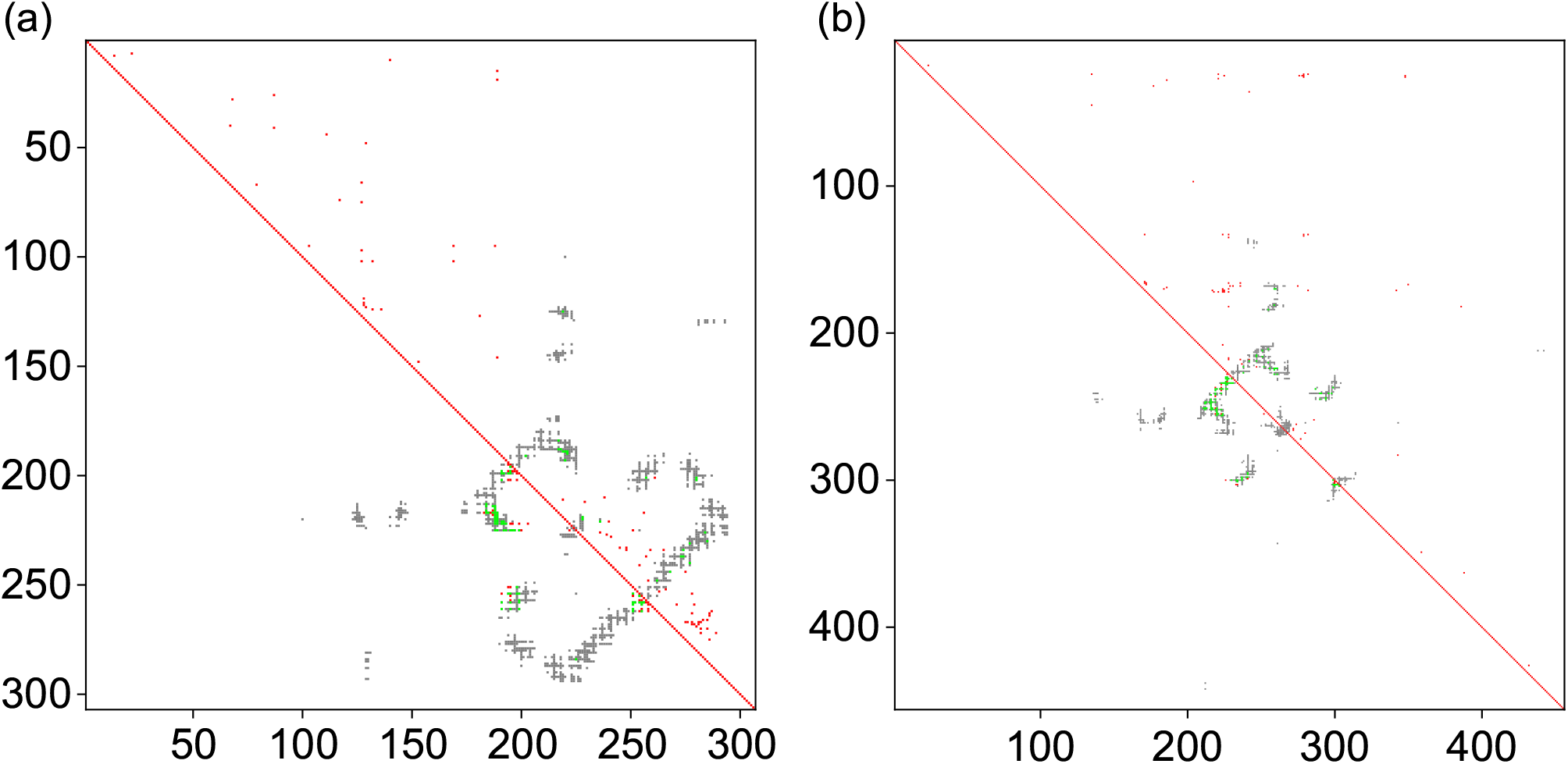
Comparison between the native contacts and the top 100 contacts predicted by DeepHomo (lower left triangle) and DCA_APC (upper right triangle) for T85 (a) and T93 (b). The native contacts, correct predictions, and incorrect predictions are colored in gray, green, and red, respectively.

**Figure 8:**
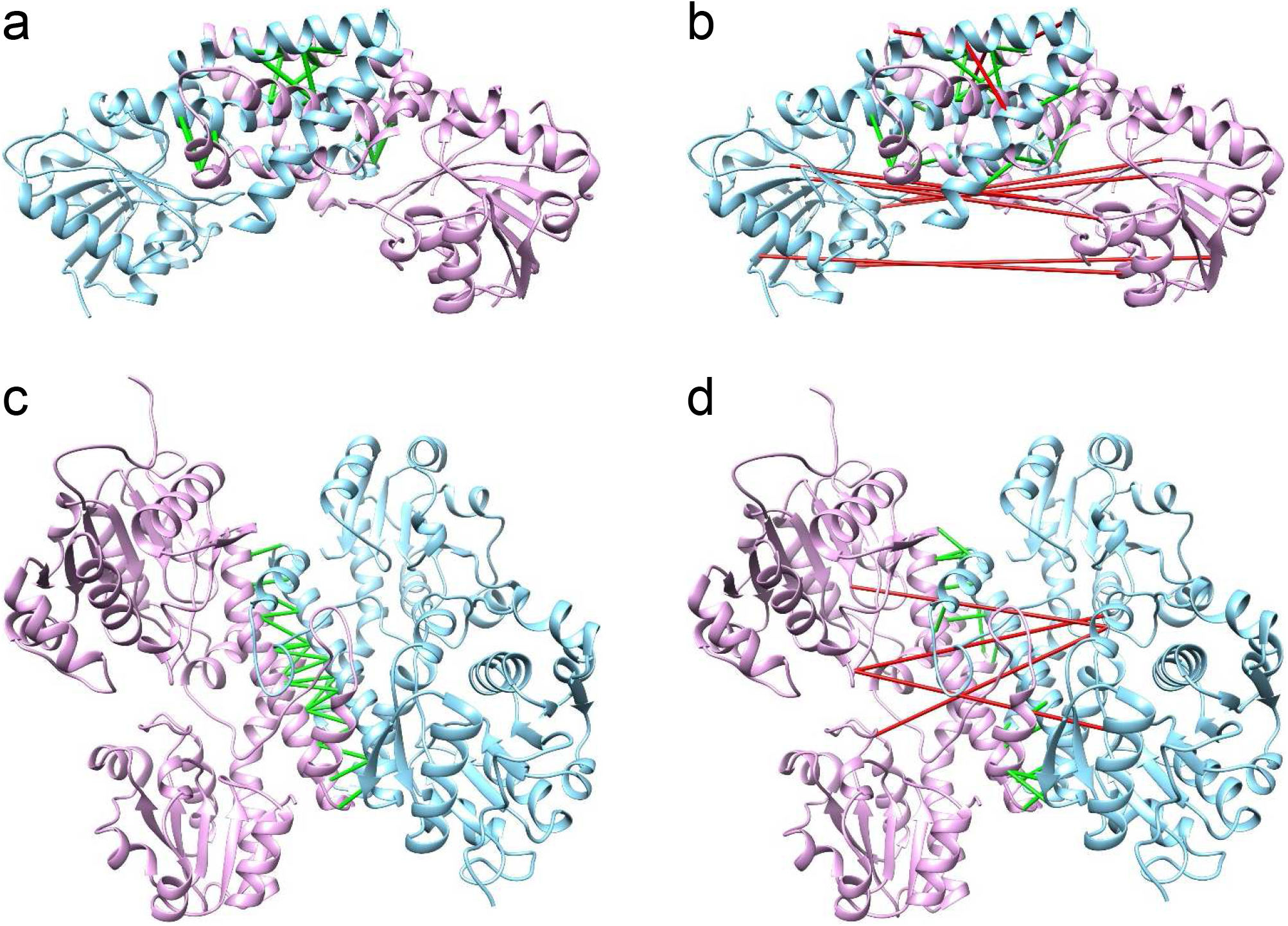
Top 10 predicted contacts by DeepHomo (a and c) and DCA_APC (b and d) on T85 (a and b) and T93 (c and d). The two monomers of native homo-dimer structures are shown in ribbons and colored in pink and blue, respectively. The correct and wrong predictions are shown in green and red connections.

### 2.4 Integration of DeepHomo into protein docking

Accurate prediction of inter-protein contacts will greatly help the structure determination of corresponding protein-protein complexes. To investigate the practical role of DeepHomo in complex structure prediction, we have integrated the predicted contacts by DeepHomo into our ab initio HSYMDOCK symmetric docking program by applying the predicted contacts in a post-docking filter, named as DeepHSYMDOCK, and evaluated its performance on the CASP-CPARI test set of 28 realistic targets. For each target, the same monomer structure, i.e. the first Zhang-Server model, was used as the input structure for the DeepHomo and HSYMDOCK processes of DeepHSYMDOCK. All the default parameters were used during the HSYMDOCK docking calculations. Given that the accuracy of contact prediction is critical to the docking process, only the top predicted contact by DeepHomo was used to filter the final binding modes predicted by HSYMDOCK.

Figure 9 shows the success rates of DeepHSYMDOCK and HSYMDOCK in binding mode prediction on the CASP-CAPRI test set. It can be seen from the figure that the predicted contacts by DeepHomo did significantly improve the docking performance in complex structure prediction, When the top five predictions are considered, DeepHSYMDOCK achieved a success rate of 64.3%, which is considerably higher than 42.9% for the ab initio HSYMDOCK docking program. Figure 10 shows a comparison between the binding modes predicted by DeepHSYMDOCK and the native structures for targets T85 and T93. Without using contacts, HSYMDOCK did not predicted any correct complex structures within the top five predictions for T85 and T93. However, with the help of the top predicted contacts. DeepHSYMDOCK achieved a medium-accuracy binding mode with *L*_rmsd_=2.804 Å, *I*_rmsd_=2.247 Å and *f*_nat_=60.211% for T85, and an acceptable-accuracy binding mode with *L*_rmsd_=7.441 Å, *I*_rmsd_=3.446 Å and *f*_nat_=29.167% for T93, within the top 2 predictions. These results demonstrated the important role of DeepHomo in the structure prediction of protein-protein complexes.

**Figure 9:**
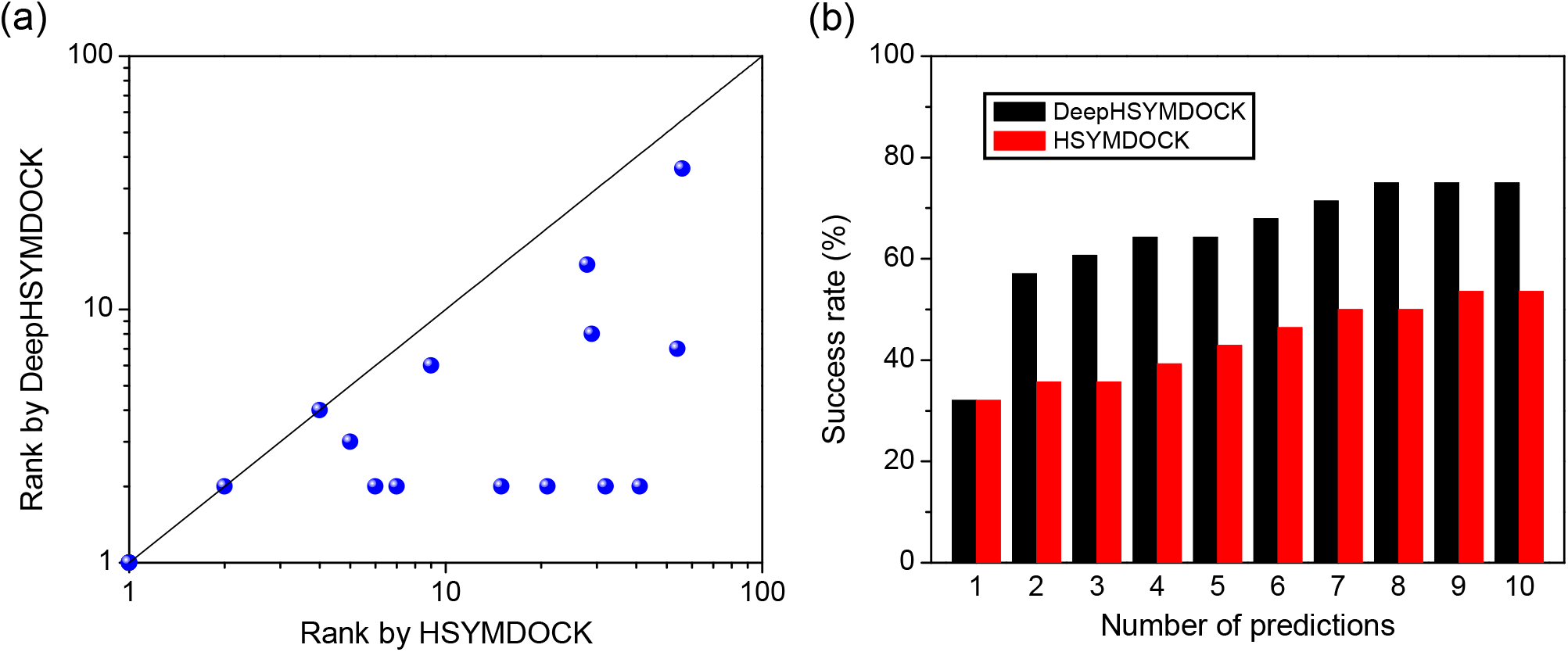
The performance of DeepHSYMDOCK, i.e. DeepHomo+HSYMDOCK, and ab initio HSYMDOCK docking algorithms in binding mode prediction on the CASP-CAPRI test set of 28 realistic targets. (a) Ranks of the first correct binding modes predicted by DeepHSYMDOCK versus those by HSYMDOCK. (b) The success rates in binding mode prediction when one to ten predictions were considered.

**Figure 10:**
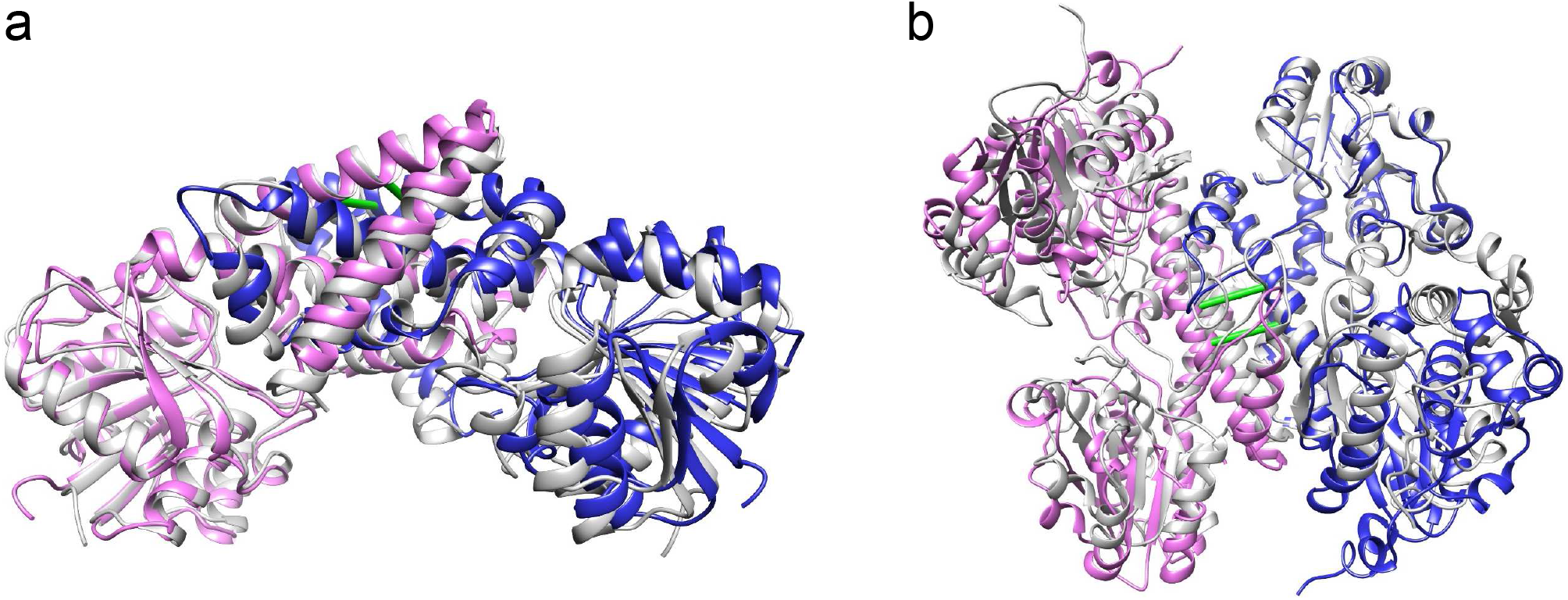
Comparison between the native structures and the predicted binding modes by DeepH-SYMDOCK. The top predicted contacts used for docking are shown in green connections. The native structure is colored in gray, and the two monomers of the predicted structure are colored in pink and blue, respectively. (a) The #2 ranked model for T85, which has a medium accuracy with *L*_rmsd_=2.804 Å, *I*_rmsd_=2.247 Å and *f*_nat_=60.211%. (a) The #2 ranked model for T93, which has an acceptable accuracy with *L*_rmsd_=7.441 Å, *I*_rmsd_=3.446 Å and *f*_nat_=29.167%.

## 3 DISCUSSIONS AND CONCLUSION

Accurate prediction of inter-protein contacts across protein-protein interfaces offer valuable structural insights into protein-protein interactions and is thus critical for the determination of corresponding complex structures. However, despite the significant progress in intra-protein contact prediction, the progress in inter-protein contact prediction remains limited because of the limited number of protein-protein complexes and the MSA joint challenge of two interacting proteins. Inspired by the great success of deep learning in recent CASP challenges for protein structure prediction, we have presented a deep learning model for inter-protein residue-residue contact prediction across homo-oligomeric protein interfaces by integrating both structural and MSA features of monomers, named as DeepHomo. Our DeepHomo model was extensively evaluated on two independent data sets, the PDB test set of 300 experimental structures and the CASP-CAPRI test set of 28 realistic homo-dimer targets, and compared with state-of-the-art DCA-based methods (DCA_APC and DCA_DI) and machine learning-based approaches (BIPSPI_seq and BIPSPI_struc). It was shown that DeepHomo achieved a high accuracy in inter-protein contact prediction and far outperformed existing DCA and machine learning-based methods. On the PDB test set, our DeepHomo obtained an accuracy of 62.7% when the top predicted contact was considered, compared with 27.0% for DCA_DI and 33.0% for DCA_APC. On the CASP-CAPRI test set, DeepHomo also obtained a high accuracy of 60.7% for the top predicted contact, which is much higher than 14.3%, 25.0%, 7.1% and 25.0% for DCA_APC, DCA_DI, BIPSPI_seq and BIPSPI_struc, respectively.

Although the monomer structure was used as an input option of our deep learning model, DeepHomo was robust to the depth of MSAs and the quality of monomer structures, and can achieve good accuracies for the MSA of ~100 sequences and the monomer structure of TM-score~0.7. The ablation analysis indicated that the complex structure of homo-dimers depended on the structure more than the sequence of monomers. Integrating the predicted contacts into protein-protein docking significantly improved the docking accuracy, suggesting the practical role of DeepHomo in the determination of homo-dimer structures. The future studies are expected to involve the inter-protein contact prediction of other homo-oligomer or hetero-oligomer complexes.

In summary, we have present a deep learning model for inter-protein contact prediction by taking homo-dimer as a start, and showed its great success on both experimentally determined and blindly predicted homo-dimeric complexes. The present model demonstrated the efficacy of integrating both structural and sequential features into a deep learning model for accurate inter-protein contact predictions. It is anticipated that DeepHomo will have a far-reaching implication in inter-protein contact prediction and sequent protein-protein complex structure determination.

## 4 MATERIALS AND METHODS

### 4.1 Deep learning architecture

As shown in Figure 1, 1D convolutional neural network is first used to capture the sequential context and extract the high-dimensional features from sequential features. Then the extracted high-dimensional sequential features are converted into pairwise features through outer concatenation as used in the RaptorX-contact study [38]. The converted features are concatenated with input pairwise features together. At last, the 2D convolutional neural network is used to capture pairwise context and output the predicted contact probability for each reside pair.

Like other deep learning methods proposed for monomer structure contact prediction, the central part of our convolutional neural network (CNN) is a stack of residual network (ResNet) blocks [57, 58]. ResNet has been widely used in computer vision and protein contact prediction because the shortcut connection added in ResNet makes the training of extremely deep CNN possible. Different with RaptorX-contact, we used ResNet v2 [58] as the basic network block for both 1D and 2D networks in this study, which has been shown to make training easier and improve generalization. The ResNet v2 block consists of two batch normalization layers, two activation layers, two convolutional layers, and a shortcut connection between the input and output of the last convolutional layer. If the input tensor of the block has a different dimension with the output one of the convolutional layer, a convolutional layer with kernel size of 1 × 1 will be used to change the dimension of the input tensor, that is the identity layer as shown in Figure 1.

The 1D CNN contains six 1D ResNet v2 blocks with an increased number of filters. A convolutional layer with two 1 × 1 filters is used to compress the output tensor of the 1D network. The kernel size of the 1D network is set to 17. The numbers of filters for the six blocks in the 1D network are set to 35, 40, 45, 50, 55 and 60, respectively. The 2D CNN is stacked by 36 2D ResNet v2 blocks with nine groups of filters. The numbers of filters for the nine groups of 2D blocks are set to 32, 32, 48, 64, 64, 64, 48, 32 and 32, respectively. The output block consists of a batch normalization layer, a activation layer, and a convolutional layer with 1 × 1 kernel where a sigmoid activation is used to convert the predicted probability in the range 0 to 1. The convolutional kernel size of the 2D network is set as 3 × 3. The exponential linear unit (ELU) is utilized as the activation function in the network. All the convolutional operations are with zero padding to maintain the size of the predicted contact map. As the training parameters in the network are independent to the size of input, our model can take variable-length proteins as input.

### 4.2 Data sets

Protein homo-oligomers have different symmetry types and stoichiometry, which may form different residue contacts at the interfaces. To avoid the effect of multiple interfaces and reduce potential noise, we focused on the homo-dimer proteins with C2 symmetry type, which is the largest class of the homo-oligomers [2]. First, we queried all the biological assemblies in the PDB with the following criteria: i) They are assigned C2 symmetry by the authors; ii) The resolution is better than 3.0 Å; iii) The biological unit only contains two protein chains; iv) The lengths of protein chains range from 50 to 500. All the queried biological assemblies were checked using blast [59] to make sure that the two chains in one assembly share more than 99% sequence identity. Then, all the candidate assemblies were clustered using MMseqs2 [60, 61] with a sequence identity cutoff of 40% to remove the redundancy. The assembly with the best resolution was chosen as the representative of the cluster. Finally, the interface area was calculated for each assembly structure and only the structures with interface more than 1000 Å^2^ were retained. This yielded a final data set of 4240 homo-dimeric complex structures, of which 3640 structures were used as the training set, 300 structures as the valid set, and 300 structures as the test set.

In addition, another independent data set, called CASP-CAPRI test set, was also constructed to evaluate the performance of our deep-learning model in realistic applications. The joint CASP-CAPRI challenge has been established since 2014, which is a double blind experiment and aims to assess the computational methods of modelling protein assemblies in the community [47–49]. We collected all the homo-dimer targets from recent four CASP-CAPRI challenges that have experimental complex structures available, yielding a total of 27 targets. In addition, we also split the CAPRI target T149 (CASP target T0999) into two interacting targets T149_D1 and T149_D4 according to the domain definition in the CASP experiment due to the large size of 1589 residues. Finally, the CASP-CAPRI test set consists of 28 homo-dimer targets with known experimental structures. To test the realistic performance of our DeepHomo model, for each target in CASP-CAPRI test set, the first Zhang-Server model, i.e. Zhang-Server-TS1, in the CASP experiments was used as the input monomer structure of DeepHomo during the evaluations.

### 4.3 Input features

In this study, two residues are defined as in contact if any two heavy atoms from the two residues are within 8 Å. Multiple features were used in our deep learning model to predict the residue contacts. These features can be grouped into two categories. One is one-dimensional (1D) sequential features including protein sequence profile like position-specific scoring matrix (PSSM), 8-state secondary structure types, and three hydrophobicity scales of each amino acid. The other is two-dimensional (2D) pairwise features, including residue-residue distance map of monomer structure, docking map created by our FFT-based docking program, and direct co-evolutionary information calculated by CCMpred [14].

Give a monomer of *L* amino acids, we first built a multiple sequence alignment (MSA) for its sequence by running HHblits [27] with a minimum coverage of 40% and a maximum pairwise sequence identity of 99% with the query sequence against the UniRef30_2020_03 database [62] by three iterations. The e-value threshold was set as the default one of 0.001. Then, according to the constructed MSA, the raw and average product correction (APC)-corrected direct coupling scores were calculated by running CCMpred. The protein sequence profile (PSSM) was also generated. The direct co-evolutionary feature was represented as a matrix with dimension *L* × *L* × 2 and the PSSM feature was represented as a matrix with size of *L* × 20.

Besides the sequence for each target, the 3D monomer structure was also used to produce the input features. It is expected that the features derived from monomer structures are helpful to distinguish inter-protein from intra-protein contacts. The DSSP (Dictionary of Protein Secondary Structure) program [63] was first used to assign the secondary structure type for each amino acid of the monomer structure. Then, the assigned secondary structure types were converted into a matrix with dimension *L* × 8 according to one-hot encoding. The ‘ACC’ column in the output of DSSP which represents the water molecules in contact with this residue was also used as a hydrophobicity feature. The absolute solvent accessible area of each residue calculated by the Naccess program [64] was used as another hydrophobicity feature. The Wimley-White whole residue hydrophobicity scale [65] was used as the third hydrophobicity feature. The pairwise distance map of the monomer structure consisted of the minimal heavy atom distances of all residue pairs with size of L × L × 1. At last, the docking map was generated from a modified version of our in-house FFT-based docking program, CHDOCK [66, 67]. The angle interval was set as 6° which resulted in 960 evenly-distributed rotations in Euler space. For each rotation, the top 100 translations according to the shape complementary score were used to generate the docking map. Specifically, for each binding mode, i.e. one predicted complex structure, a contact map for the interaction of two chains was constructed with the same definition as described above. Then, all the 96000 contact maps were combined into one docking map with size of *L* × *L* × 1. For each target in the training, valid, and test set, the monomer structure was directly exacted from the experimental complex structure. For each target in the CASP-CAPRI test set, the top Zhang-Server model from CASP was used as the monomer structure.

### 4.4 Implementation

The Keras with the Tensorflow as backend was used to implement our deep learning model. The hyper-parameters were set as follows: mini-batch size: 1, optimizer: Adam [68], learning rate: 0.001, and L_2_-norm regularization coefficient: 1e-4. The L_2_-norm regularization was applied by adding a weight decay to each convolutional layer. The learning rate decay strategy was set as follows. If the loss of the valid set stops improving in two epochs, the learning rate will reduce by a factor of 0.2 and the minimal learning rate was set as 1e-6. Due to the extremely unbalance between non-contact residue pairs and the contact ones, here we used the Focal Loss [46] as the loss function to train our deep learning model. The loss function for each residue pair of *i* and *j* is described as follow.

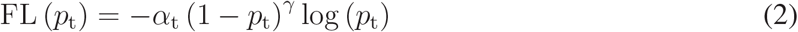

where *α*_*t*_ and *p*_*t*_ are defined as

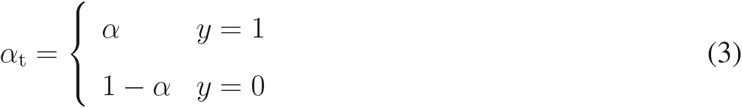

and

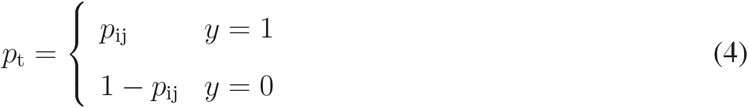

Here, *p*_*ij*_ is the predicted contact score of reside *i* and residue *j*, *α* is the parameter to balance the importance of positive and negative examples, and *γ* is the tunable focusing parameter to focus learning on hard examples and down-weight the numerous easy examples. In our model, *α* is set as 0.25 and *γ* was set to 2, which are the same those in the original paper. Due to the limited graphics processing unit (GPU) memory, the maximum length of the protein fed into the network was limited to 400. If the length is more than 400, a continuous subsequence with length of 400 is randomly sampled to represent the protein. The model was trained on one K80 GPU and after around 30 epochs (about 40h), the model converged to a stable solution.

As homo-dimer protein often has a C2 symmetry type, the ideal contact map for a homo-dimer interface is diagonally symmetric. Hence, the labels and the input features used to train our model have to be operated to ensure the symmetry. For the input pairwise features, the transposed and untransposed matrixes were averaged to get the final input symmetric matrix. For the labels of the contact map, if the residue pair (*i, j*) was with label of 1, the residue pair (*j, i*) would be set to 1 too, and vice versa. Mask records were created for missing residue pairs in the complex structure to ignore those missing residue pairs in the loss function.

### 4.5 Evaluation criteria

The performance of residue-residue contact prediction was measured by two parameters. One is the accuracy, which is defined as the percentage of true positive (TP) contacts among the top *n* predicted contacts. As a few correct restraints may significantly help filter the correct binding modes for molecular docking [69], so the accuracy of the top contact predictions is crucial for complex structure prediction. The average accuracy of all the targets was used to represent the performance of a model on the data set. The other is the success rate, which is defined the percentage of the homo-dimer targets with at least one successfully predicted contact when a certain number of predicted contacts are considered, compared to all the targets in the test set. Such success rate parameter is useful if users want to validate the predicted contacts using experimental methods, because they will know how many pairs of residues need to be examined.

For docking applications, the quality of a predicted binding mode is measured by the CAPRI criteria and were be divided into four categories: high, medium, acceptable, and incorrect [70]. The success rate is used to measure the docking performance, which is defined as the percentage of the targets with at least one successful (acceptable or better accuracy) prediction among the total number of targets in the test set when a certain number of top predictions are considered.

## ACKNOWLEDGEMENTS

This work is supported by the National Natural Science Foundation n 2016YFC1305805), and the startup grant of Huazhong University of Science and Technology.

## Author contributions

S.H. conceived and supervised the project. Y.Y. and S.H. designed and performed the experiments. Y.Y. and S.H. wrote the manuscript.

## Competing interests

The authors declare no competing interests.

## Code availability

DeepHomo is freely available for academic use through http://huanglab.phys.hust.edu.cn/DeepHomo/.

## Notes

### Competing Interest Statement

The authors have declared no competing interest.

